# Harmonizing Heterogeneous Single-cell Gene Expression Data with Individual-level Covariate Information

**DOI:** 10.1101/2025.04.15.649009

**Authors:** Yudi Mu, Wei Vivian Li

**Affiliations:** Department of Statistics, University of California Riverside, Riverside, CA 92521, USA

## Abstract

**Motivation:** The growing availability of single-cell RNA sequencing (scRNA-seq) data highlights the necessity for robust integration methods to uncover both shared and unique cellular features across samples. These datasets often exhibit technical variations and biological differences, complicating integrative analyses. While numerous integration methods have been proposed, many fail to account for individual-level covariates or are limited to discrete variables.

**Results:** To address these limitations, we propose scINSIGHT2, a generalized linear latent variable model that accommodates both continuous covariates, such as age, and discrete factors, such as disease conditions. Through both simulation studies and real-data applications, we demonstrate that scINSIGHT2 accurately harmonizes scRNA-seq datasets, whether from single or multiple sources. These results highlight scINSIGHT2’s utility in capturing meaningful biological insights from scRNA-seq data while accounting for individual-level variation.

**Availability and implementation:** The scINSIGHT2 method has been implemented as an R package, which is available at https://github.com/yudimu/scINSIGHT2/.

## 1 Introduction

Single cell RNA sequencing (scRNA-seq) technologies have transformed our understanding of cellular diversity and gene expression dynamics [1, 2, 3]. They have also enriched our insights into biological processes and disease mechanisms by profiling transcriptomes of individual cells [4, 5]. However, the analysis of scRNA-seq data poses significant challenges due to both biological and technical variations, including differences in experimental protocols, sequencing depth, and subject-level differences represented by covariates like gender, age, and disease status [6, 7, 8]. Such variations can lead to inconsistent gene expression patterns that obscure genuine cellular features and hinder comparisons across studies. Therefore, effective integration of multiple scRNA-seq samples is crucial for accurately clustering cells, annotating them, and interpreting their functions, thereby enhancing our understanding of cellular identities.

A variety of integration methods exist for scRNA-seq data. Many of them perform integration based on cells with mutually similar expression profiles between different single-cell samples, which are termed mutual nearest neighbors (MNNs). For example, mnnCorrect [9] initially identifies MNNs between single-cell samples and then attempts to enhance data consistency by removing the differences between MNN cells, assuming these differences only represent technical variations. Other methods have refined this approach in different ways. Seurat [10] identifies MNNs within a shared low-dimensional space learned via canonical correlation analysis; Scanorama [11] searches for MNNs after conducting dimensionality reduction through randomized singular value decomposition; and Harmony [12] introduces a soft clustering process, opting to perform data correction within clusters rather than strictly between MNNs. In addition, some methods employ matrix factorization to elucidate both shared and sample-specific factors, thereby facilitating the identification of cell populations across samples. Example methods include LIGER [13], CSMF [14], and scCoGAPS [15]. Moreover, generative models such as autoencoders and variational autoencoders, employed by tools like scGen [16], scVI [17], and DESC [18], create a unified embedded space to integrate single-cell samples.

While most existing scRNA-seq integration methods primarily aim to mitigate technical variations, often referred to as batch effects, a few methods have recently been developed to explicitly address biological differences among single-cell samples from diverse conditions, such as varying experimental groups or disease conditions. For instance, our previous work, scINSIGHT [19], leverages a non-negative matrix factorization model to discern coordinated gene expression patterns that are either common or unique to different biological conditions. Similarly, scDisco [20] utilizes a variational autoencoder to integrate condition factors in joint analysis of single-cell samples. Another autoencoder-based approach, scDisInfact [21], employs multiple encoders to model more than one discrete condition factors.

Despite these advancements, current methods concentrate solely on distinct types of biological variations like gender or disease status, which restricts their ability to harness the full spectrum of covariate data, particularly continuous biological variables such as age. To bridge this gap, we introduce scINSIGHT2 (Figure 1), a new integration model designed to harmonize gene expression data from multiple single-cell samples by incorporating both discrete and continuous individual-level covariates. Utilizing a generalized linear latent variable model, scINSIGHT2 adjusts for covariate-associated gene expression changes prior to estimating cell embeddings within a unified low-dimensional space of inferred metagenes. Our evaluations through both simulations and real-data applications have demonstrated that the latent cell embeddings and metagenes identified by scINSIGHT2 effectively capture meaningful cellular identities. This capability allows scINSIGHT2 to enhance the annotation and interpretation of cell populations across diverse single-cell samples.

**Figure 1:**
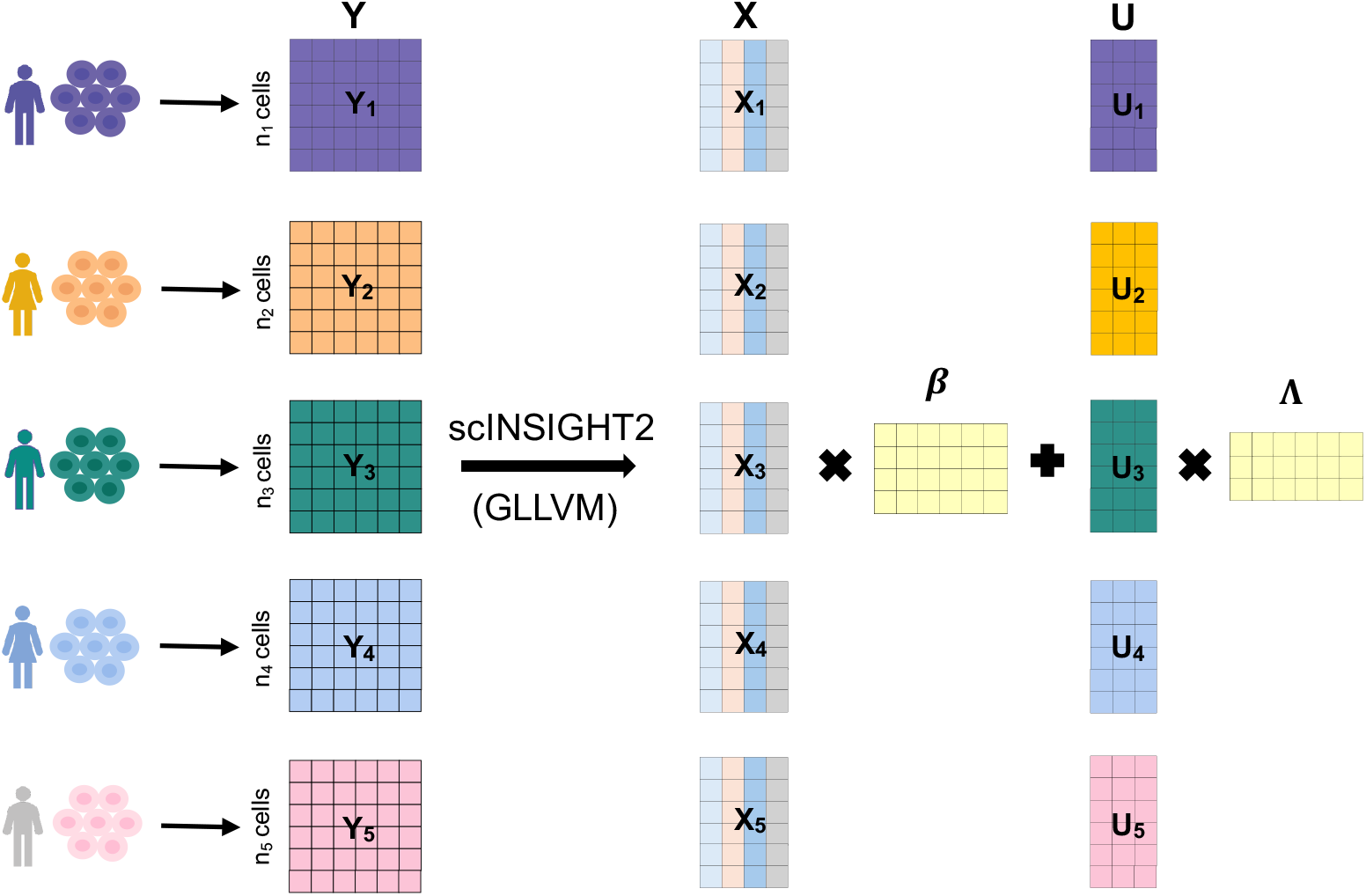
An overview of the scINSIGHT2 method. In this toy example, we assume five single-cell samples (*Y*_1_, *…, Y*_5_) from different subjects. In GLLVM, the samples are concatenated together as a large count matrix *Y*. Matrix *X* represents the covariate information, where each column represents a different covariate such as age, gender or BMI. The output of the scINSIGHT2 model consists of the inferred cell embeddings *U* and the corresponding loading matrix of the metagenes Λ. Then, the joint clustering analysis is performed on *U* to obtain the cellular identities.

## 2 Methods

### The scINSIGHT2 model

We assume there are *K* scRNA-seq samples from *K* subjects, each summarized as a count matrix *Y*_*k*_ (*k* = 1, …, *K*). After gene and cell filtering, these count matrices have *M* common genes, and sample *k* has *n*_*k*_ cells. We use *Y* to denote the concatenated count matrix of all *K* samples, which includes 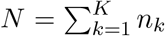 cells and *M* genes. In addition to single-cell gene expression data, we assume the availability of *D* covariates at the subject level, such as age, gender, and clinical variables. By explicitly modeling these variables and batch information in scINSIGHT2, our method attempts to remove batch effects in gene expression data that are introduced by these factors.

In scINSIGHT2, we construct a generalized linear latent variable model (GLLVM) to enable the joint dimensionality reduction of multiple single-cell samples while accounting for subject-level covariates (Figure 1). For each element in matrix *Y*, we assume *y*_*ij*_ ∼ Poisson(*μ*_*ij*_), and its mean parameter *μ*_*ij*_ is modeled via a log link function:

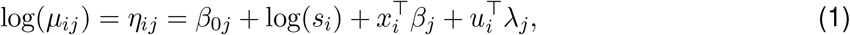

where *y*_*ij*_ is the count of gene *j* (*j* = 1, …, *M*) in cell *i* (*i* = 1, …, *N*); *x*_*i*_ ∈ ℝ^*D*^ denotes the covariates of the sample that the *i*th cell belongs to; *s*_*i*_ is the library size factor for cell *i*. We define the library size for cell *i* as 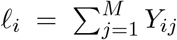, and the library size factor is calculated as *s*_*i*_ = *l*_*i*_*/*median{*l*_*i*_|*i* = 1, …, *N*}. Among the unknown parameters or latent factors, *β*_0*j*_ is the intercept; *β*_*j*_ ∈ ℝ^*D*^ denotes gene-specific regression coefficients corresponding to the covariates; *u*_*i*_ ∈ ℝ^*P*^ (*P* ≪ *M*) denotes latent embeddings of cell *i*, which represent the cell’s scores on latent metagenes (linear combinations of the original gene features); *λ*_*j*_ ∈ ℝ^*P*^ denotes the loadings of gene *j* on the *P* metagenes.

In order to properly estimate the unknown parameters, we make three assumptions. (1) The cell embeddings are *U* = [*u*_1_, …, *u*_*P*_], and each element *u*_*i*_ ∼ 𝒩 0, *σ*^2^*I*_*P*_, where *σ*^2^ is the variance to be estimated; (2) Λ = [*λ*_1_, *λ*_2_, …, *λ*_*M*_] ∈ ℝ^*P* ×*M*^ is a lower triangular matrix with positive elements on the diagonal; (3) Conditioned on *u*_*i*_, the responses *y*_*ij*_ are independent of each other. Assumptions (1) and (2) are used to ensure parameter identifiability [22]; otherwise, *u*_*i*_ and *λ*_*j*_ could be rotated without affecting the value of their product. It is important to note that these assumptions do not compromise model performance, as our primary downstream analysis involves clustering cells based on *u*_*i*_, where the relative positions of the cells embedded in *u*_*i*_ remain unchanged. Assumption (3) is common in GLLVMs in order to derive the likelihood function as follows.

We use *f* (*y*_*ij*_ | *u*_*i*_, Θ) to denote the conditional density function of *y*_*ij*_ given the cell embed-dings, where 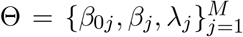 denote the model parameters to be estimated. The marginal log-likelihood function after integrating out the cell embeddings is:

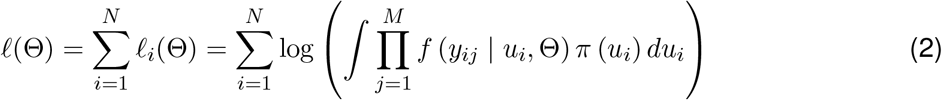

where *ℓ*_*i*_(Θ) is the marginal log-likelihood of cell *i*; *π* (*u*_*i*_) ∼ 𝒩 0, *σ*^2^*I*_*P*_, as stated in Assumption (1). For *ℓ*_*i*_(Θ), we have:

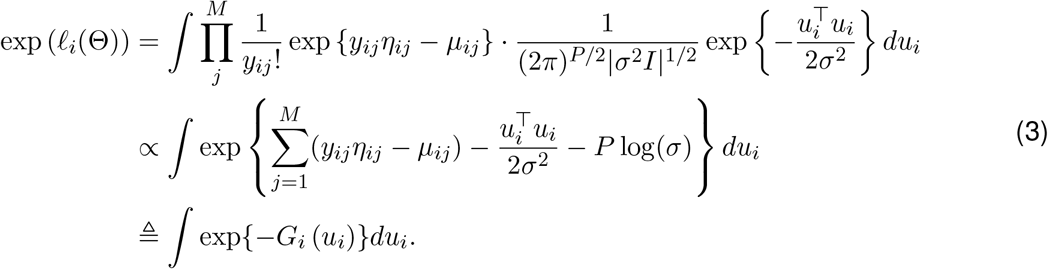

Since the integral part does not have a closed form solution, we use the Laplace method to approximate the log-likelihood function as proposed in [23]. According to the Laplace method [24], we can approximate *ℓ*_*i*_(Θ) as

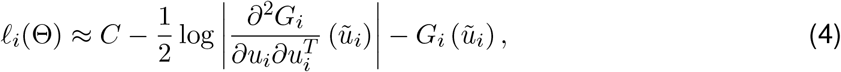

where 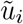 is the solution to *∂G*_*i*_(*u*)*/∂u*_*i*_ = 0 (i.e., 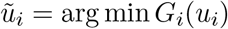), and *C* is a constant that does not depend on 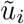. Given the derivatives 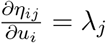 and 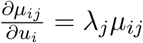, we have

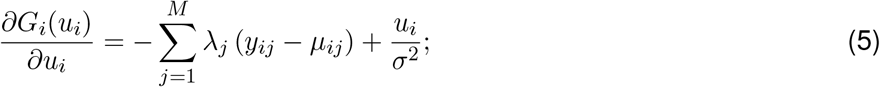

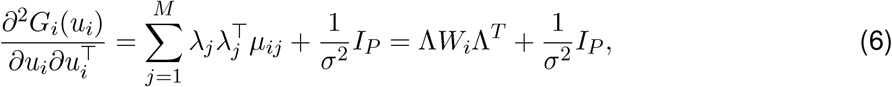

where *W*_*i*_ is a diagonal matrix with [*W*_*i*_]_*jj*_ = *μ*_*ij*_ for *j* = 1, …, *M*. Then, equation (4) becomes

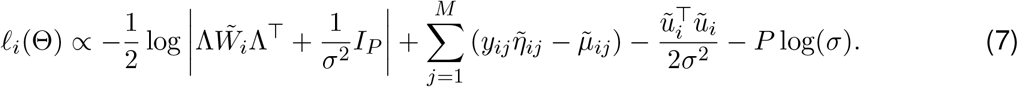

When *M* and *N* are large, which is usually the case with scRNA-seq data, the first term in equation (7) is asymptotically dominated by the second term [25]. Therefore, we will ignore the first term when we optimize the likehood function. Then, we could define the objective function below to approximate the maximum likelihood estimation problem:

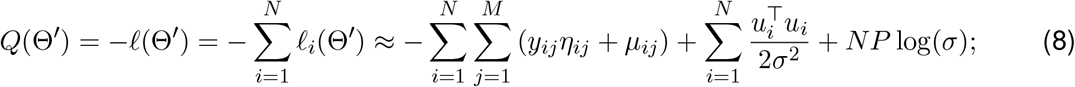

where 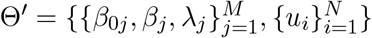. Finally, we can obtain the estimators 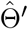 using the gradient descent method (see Supplementary Methods).

Once all the parameters have been estimated, we use the estimated cell embeddings *Û*_*N* ×*P*_ to perform joint clustering analysis across multiple single-cell samples. As the batch effects introduced by covariates have been removed during the modeling process, these cell embeddings are expected to reveal cellular identities that might have been obscured in the original gene expression data. However, our estimation process relies on the Laplace approximation and may not directly yield cell embeddings that are optimal for clustering. Therefore, we implement a normalization procedure from the scINSIGHT method [19] to further refine the cell embeddings before applying the Louvain method [26] to identify cell populations (see Supplementary Methods).

### Selection of metagene number

Before performing the estimation procedure, we need to determine the value of *P*, which represents the number of metagenes or latent factors in GLLVM. Although users can run the model with different values of *P* and investigate the corresponding results in an interactive manner, we provide a heuristic approach here to guide the selection of this hyperparameter.

Denote the candidate set of values (ordered from the smallest to the largest) as *A*. For each candidate value *p* ∈ *A*, we run our model *B* (defaults to 10) times with different seeds in the initialization step. After obtaining the clustering results for the *b*-th (*b* = 1, …, *B*) initialization, we construct a connectivity matrix *C*^(*p,b*)^ of size *N* × *N* between the cells. In this matrix, 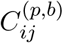 is set to 1 if cell *i* and cell *j* belong to the same cluster, and 0 if they belong to different clusters. Then, the consensus matrix is defined as 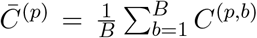. Based on the consensus matrix, the corresponding consensus index is calculated, defined as

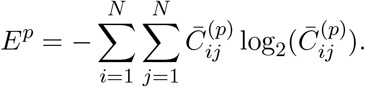

If 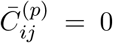 the product term is set to 0. We expect the consensus index to be close to 0 if the clustering results across different initializations are highly consistent; in contrast, the index will have a large value if the results across different initializations are very different. Once we have calculated the consensus indices for all candidate parameters, we identify a subset *A*^0^ ⊆ *A* consisting of candidate values whose corresponding consensus indices are lower than those of their preceding and succeeding candidates. We then select the smallest value in *A*^0^ as the metagene number. This approach seeks a metagene number that yields stable clustering results, while also favoring more parsimonious models.

### Simulation study

We generated synthetic single-cell gene expression data in different settings to evaluate the performance of scINSIGHT2. In the first setting, we generated data for 10 subjects, each with 500 cells and 2000 genes. In addition, we considered five covariates: age, gender (female or male), disease condition (healthy, mild or severe), body mass index (BMI) and subject ID. For each subject, their age was randomly simulated in the range between 20 and 70; gender and disease condition were randomly chosen; BMI was randomly simulated in the range between 16 and 24. Both age and BMI values were standardized to have a mean of 0 and a standard deviation of 1.

For the gene expression data, the simulation procedure included the following major steps. (1) We simulated library sizes based on a COVID-19 scRNA-seq dataset [27]. We selected the B cells in this dataset, and only retained genes that were expressed in a minimum of 3 cells and cells with at least 200 expressed genes. Subsequently, we utilized the Seurat package [10] to identify the top 2000 highly variable genes. Following these procedures, we obtained a count matrix consisting of 1639 cells and 2000 genes, from which we calculated the library sizes, and then sampled 5000 values to be used for generating the simulated data. (2) For the regression coefficients, the intercept *β*_0*j*_ was randomly simulated from a standard normal distribution. For each of the covariates among age, gender and BMI, we randomly selected 300 genes that were impacted by each of these factors, and the other 1700 genes were independent of these factors. For the 300 genes, their corresponding regression coefficients were randomly drawn from Uniform(1, 1.5) or Uniform(−1.5, −1) with equal probabilities; for the other 1700 genes, the regression coefficients were set to 0. For the disease condition, the regression coefficients for all genes were set to 0. This was done to incorporate a covariate not affecting gene expression, as might occur in practical applications. (3) To simulate the true cell embeddings, we assumed that there were 8 cell types, each accounting for 12.5% cells in the samples. The true number of metagenes was set to 5. Each element in *U* followed a normal distribution with a standard deviation of 0.1. The mean of the normal distribution varied for different cell types and metagenes. Supplementary Table S1 summarized the mean parameters for different cell types and metagenes. When simulating the cell embeddings, we deviated from Assumption (1) in the model. This deviation allowed us to confirm that violating this assumption does not compromise the performance of scINSIGHT2. (4) In order to generate the loading matrix Λ for the metagenes, we estimate the information from the real data mentioned above. We applied scINSIGHT2 to estimate the loading matrix, accounting for the age and gender covariates. The estimated matrix was then treated as the ground truth matrix in this simulation. (5) After simulating all the above components, we calculated 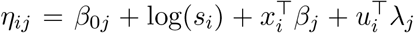. Subsequently, we could simulate the count data *Y*, where for each cell *i, Y*_*i*·_ followed a multinomial distribution with a sample size corresponding to the library size *s*_*i*_ and the probability of gene *j* being 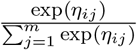.

In the second simulation setting, we explored scenarios where certain cell types were unique to specific biological conditions. Specifically, we assumed that cell type 1 was exclusive to healthy condition, and cell type 2 was exclusive to mild condition, with all other settings remaining the same. Under healthy condition, the proportions of cell types were 1*/*7 for each, excluding the cell type 2 which was 0. Under mild condition, the proportions were 0 for cell type 1 and 1*/*7 for the remaining cell types. For severe condition, cell types 1 and 2 were absent, and the remaining cell types each had a proportion of 1*/*6.

In the third simulation setting, we assumed that the proportions of two cell types (cell types 1 and 2) were dependent on age. For subjects under the age of 30, the proportions of the 8 cell types were: 0.3 for cell type 1, and 0.1 for the remaining cell types. For those aged 30 to 44, the proportions were: 0.2 each for cell types 1 and 2, and 0.1 for the remaining cell types. For subjects aged 45 and above, the proportions were: 0.3 for cell type 2, and 0.1 for the remaining cell types.

### Covariate-specific integration index

We extended the integration score used in scINSIGHT [19] so that it can be applied to both continuous and categorical variables. We define an integration index to quantify how well the integrated data removes sample-specific effects with respect to each covariate. After we get the cell embeddings from scINSIGHT2 or an alternative method, we find two sets of nearest neighbors for each cell *i* (*i* = 1, …, *N*). The first set of nearest neighbors contains *k*_1_ = 50 cells, and we denote this set as *N*_1*i*_. The second set contains *k*_2_ = 500 nearest neighbors, and we denote it as *N*_2*i*_. Then, for the *d*-th covariate, the covariate-specific integration index is defined as:

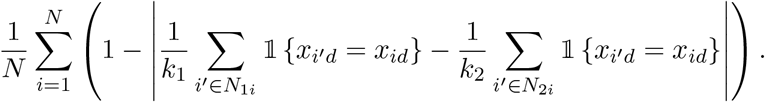

The integration index ranges from 0 to 1. It measures how frequently cells with the same covariate value appear in a local neighborhood compared to their frequency in a broader population. A higher integration index suggests that cells from subjects with varying covariate values are well integrated.

### Implementation of alternative methods

The Seurat, Harmony, scINSIGHT, scDisInFact, Scanorama, and scVI methods were applied using R package `Seurat` (v4.4.0), R package `harmony` (v1.0.3), R package `scINSIGHT` (v0.1.4), Python package `scDisInFact` (v0.1.0), Python package `scanorama` (v1.7.4), and Python package `scvi-tools` (v1.1.5), respectively. Seurat, Scanorama, and scVI do not take any additional covariate information; therefore, we used their default analysis pipelines. scINSIGHT is limited to handling a single biological covariate. In our simulation study, we used age groups as the input covariate. Harmony and scDisInFact are able to take categorical covariate information. In all analyses, categorical variables like gender and subject ID were directly input to Harmony and scDisInFact; for other continuous variables, we first categorized those variables into non-overlapping groups before used them as inputs. To cluster observed data, we directly input the full count matrix into the Seurat clustering pipeline.

### Data analysis

In both simulation and real data studies, before performing integration with different methods, we first identified the top 2000 highly variable genes using the FindVariableFeatures function of the Seurat package. In all simulation studies, the candidate set of metagene number *P* was {3, 4, …, 10}. The number of metagene numbers selected by scINSIGHT2 was 4, 4, and 3 in the three simulation settings, respectively. For Harmony and scDisInFact, we first categorized age into 3 groups: age *<* 30, 30 ≤ age *<* 45, and age ≥ 45, and BMI into 2 groups: BMI ≤ 18 and BMI *>* 18.

In the analysis of pancreatic data, we selected common cell types in the two real datasets [28, 29]: acinar, alpha, beta, delta, ductal, endothelial, epsilon, and macrophage cells. scINSIGHT2 selected the metagene number *P* as 20 from candidate values in {5, 10, 15, 20, 25, 30}. For Harmony and scDisInFact, the age variable was divided into three groups to make it a discrete variable: age *<* 30, 30 ≤ age *<* 50 and age ≥ 50. Similarly, BMI was divided into three groups: BMI *<* 22, 22 ≤ BMI *<* 29 and BMI ≥ 29. In the analysis of acinar cells, differential gene expression (DGE) analysis was performed using the Wilcoxon test. Only genes that were detected in a minimum fraction of 0.8 in either cluster were tested. The differentially expressed genes (DEGs) were those genes such that the false discovery rate (FDR) was less than 0.05, and the absolute log fold change was greater than 0.4. In order to compare the marker genes of clusters generated by different methods, we performed DGE analysis using the zlm function in the MAST package [30], where age and gender were included as covariates. The DEGs were those genes such that the FDR was less than 0.05, and the absolute log fold change was greater than 1.5.

In the analysis of COVID-19 data [31], scINSIGHT2 picked the metagene number *P* as 25 from candidate values in {5, 10, 15, 20, 25, 30}. For Harmony and scDisInFact, age was divided to three groups: age *<* 35, 35 ≤ age *<* 55 and age ≥ 55. To identify DEGs, we used the zlm function in MAST package, with age, gender, and disease condition as covariates. The gene ontology (GO) enrichment analysis was performed using the clusterProfiler package [32].

## 3 Results

## 3.1 Simulation

To assess the performance of our model, we generated single-cell gene expression data with predefined cell identities and subject-level covariates (see Methods for details). We created ten single-cell samples, each corresponding to a different subject and containing eight distinct cell types. To follow real-world scenarios, we incorporated three covariates—age, gender, and BMI—that influenced the expression levels of certain genes. Additionally, we included a categorical disease condition variable as a covariate unassociated with gene expression. An ideal integration method should consistently identify cellular identities across subjects, seamlessly accounting for inter-individual variations due to these covariates. In addition to scINSIGHT2, we compared the performance of five alternative methods: Seurat, Harmony, Scanorama, scVI, and scDisInFact. The first four methods have demonstrated strong performance in previous benchmark studies [33, 34, 35], and scDisInFact is a more recent method that can model categorical covariates.

We applied the six integration methods to the simulated data, generating low-dimensional cell embeddings and computationally inferred cell clusters as depicted in Figure 2A. The results suggest that scINSIGHT2 more accurately clustered cells than the alternative methods. Notably, it effectively mitigated the batch effects associated with age, gender, and BMI (Supplementary Figure S1), enhancing its clustering precision. In contrast, the cell distributions from Seurat, Harmony, scDisInFact, and Scanorama indicate that their embeddings still exhibited residual batch effects after integration. We compared the clustering performance of scINSIGHT2 and the five alternative methods based on the adjusted Rand index (ARI), whose higher value indicates a better agreement between the true cell types and inferred cluster labels. scINSIGHT2 achieved the highest ARI score among all the methods (Figure 2B), followed by scDisInFact and Seurat. To quantitatively evaluate methods’ effectiveness of integrating samples and removing covariate-specific effects, we introduced the covariate-specific integration index (see Methods). This index ranges between 0 and 1, with a higher value indicating that cells from subjects with varying covariate values are well integrated. scINSIGHT2 achieved the highest average integration index across varying covariates, followed by scVI and Seurat (Figure 2C). To analyze whether the inclusion of continuous covariates enhances integration in scINSIGHT2, we tested the model on the same simulated data but omitted the two continuous covariates: age and BMI. The ARI score decreased from 0.918 to 0.783, indicating a reduction in clustering accuracy. Despite this decrease, the score remained higher than those achieved by Harmony and Scanorama. These findings confirms the beneficial impact of accounting for continuous covariates in scRNA-seq integration problems when such information is available.

**Figure 2:**
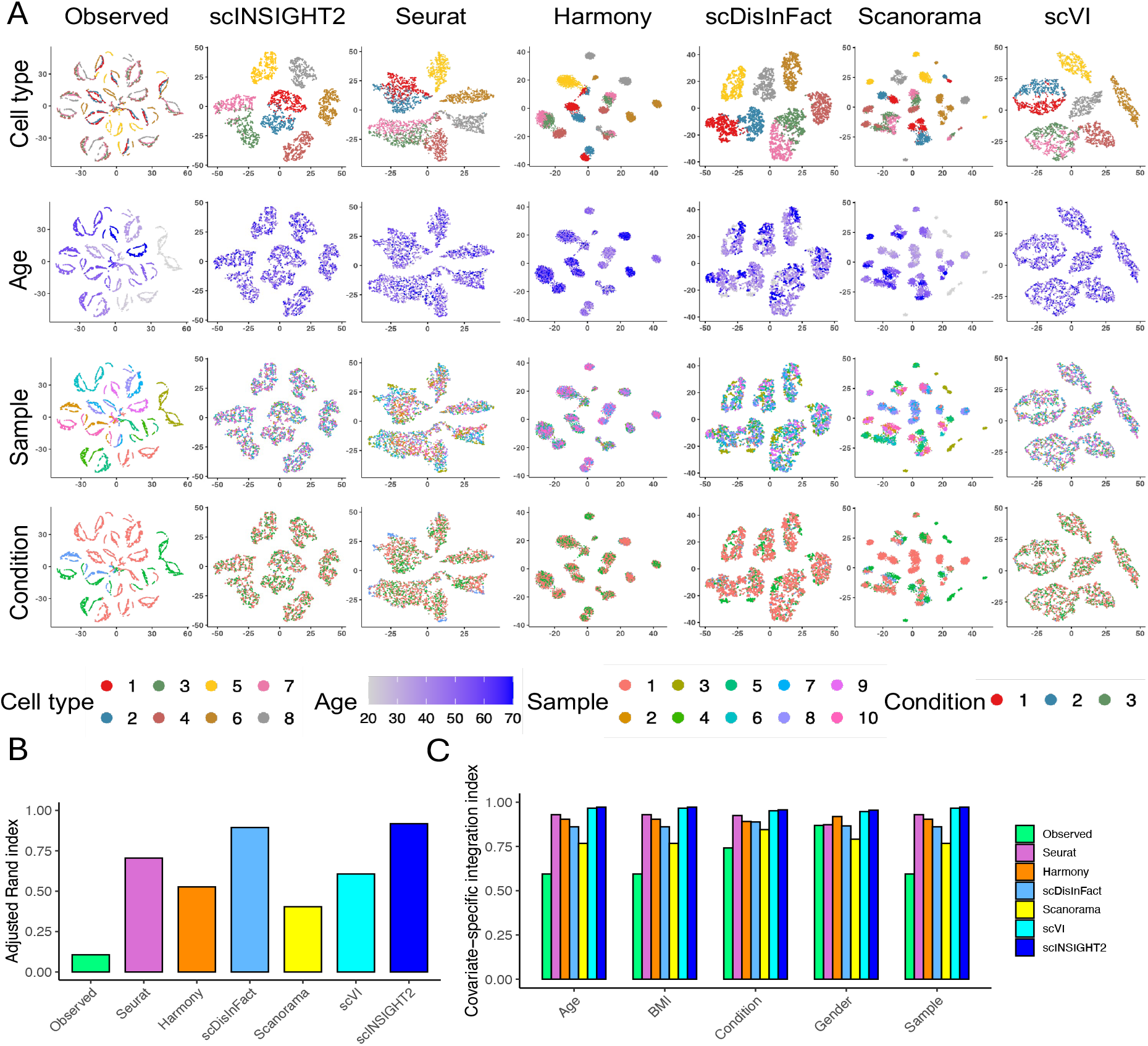
Comparison between integration methods in the simulation study (under first setting). (**A**) tSNE plots of observed data and integrated data by different methods. For each method, four tSNE plots colored by true cell type, age, sample ID, and condition are displayed. (**B**) Adjusted Rand index calculated using clusters identified from the observed or integrated data. (**C**) Covariate-specific integration index of the observed and integrated data.

Additionally, we compared scINSIGHT2 with our previous method scINSIGHT, which was designed to integrate scRNA-seq data across varying biological conditions. Given that scINSIGHT can only incorporate a single categorical covariate, it is not anticipated to surpass scINSIGHT2 in this simulation. As expected, scINSIGHT achieved an adjusted Rand index (ARI) of only 0.518. It was unable to effectively eliminate the batch effects caused by age, BMI, and gender (Supplementary Figure S2).

In addition to the above simulation, we considered two modified scenarios to more comprehensively compare different methods. First, we considered the presence of unique cell types in different biological conditions. Specifically, cell types 1 and 2 only existed in conditions 1 and 2, respectively. In this scenario, we still observed that scINSIGHT2 and scDisInFact yielded the highest ARI scores, while scINSIGHT2 and scVI achieved the best covariate-specific integration indices (Supplementary Figure S3). Second, we evaluated a scenario where the abundance of cell types 1 and 2 varied with age. Here, the proportion of cell type 1 increased with age, while that of cell type 2 decreased. Similarly, scINSIGHT2 and scDisInFact yielded the best ARI scores (Supplementary Figure S4), with scINSIGHT2 performing better in removing the covariate effects.

### 3.2 Application to human pancreas data

After evaluating the methods on simulated data, we extended our analysis to a cross-study application using single-cell samples from human pancreases. We conducted a joint analysis of 15 single-cell samples derived from two distinct studies: four donors from a 2016 study [29] and 11 donors from a 2022 study [28]. Following preprocessing (see Methods), the dataset comprised 27, 435 cells, representing eight distinct cell types. The covariates available for this analysis were age and gender.

We applied scINSIGHT2 and the five alternativie methods to this dataset to obtain cell embeddings and cell cluster labels. scINSIGHT2 identified 10 clusters based on its estimated latent factors, while Seurat, Harmony, scDisInFact, Scanorama, and scVI identified 18, 17, 23, 28, and 22 clusters, respectively (Supplementary Figure S5). We then compared these clusters with the annotated cell types from the original publications based on the ARI score. scVI and scINSIGHT2 achieved the highest ARI scores among all methods (Figure 3A). Additionally, scINSIGHT2 and Seurat achieved the highest covariate-specific integration indices (Figure 3B).

**Figure 3:**
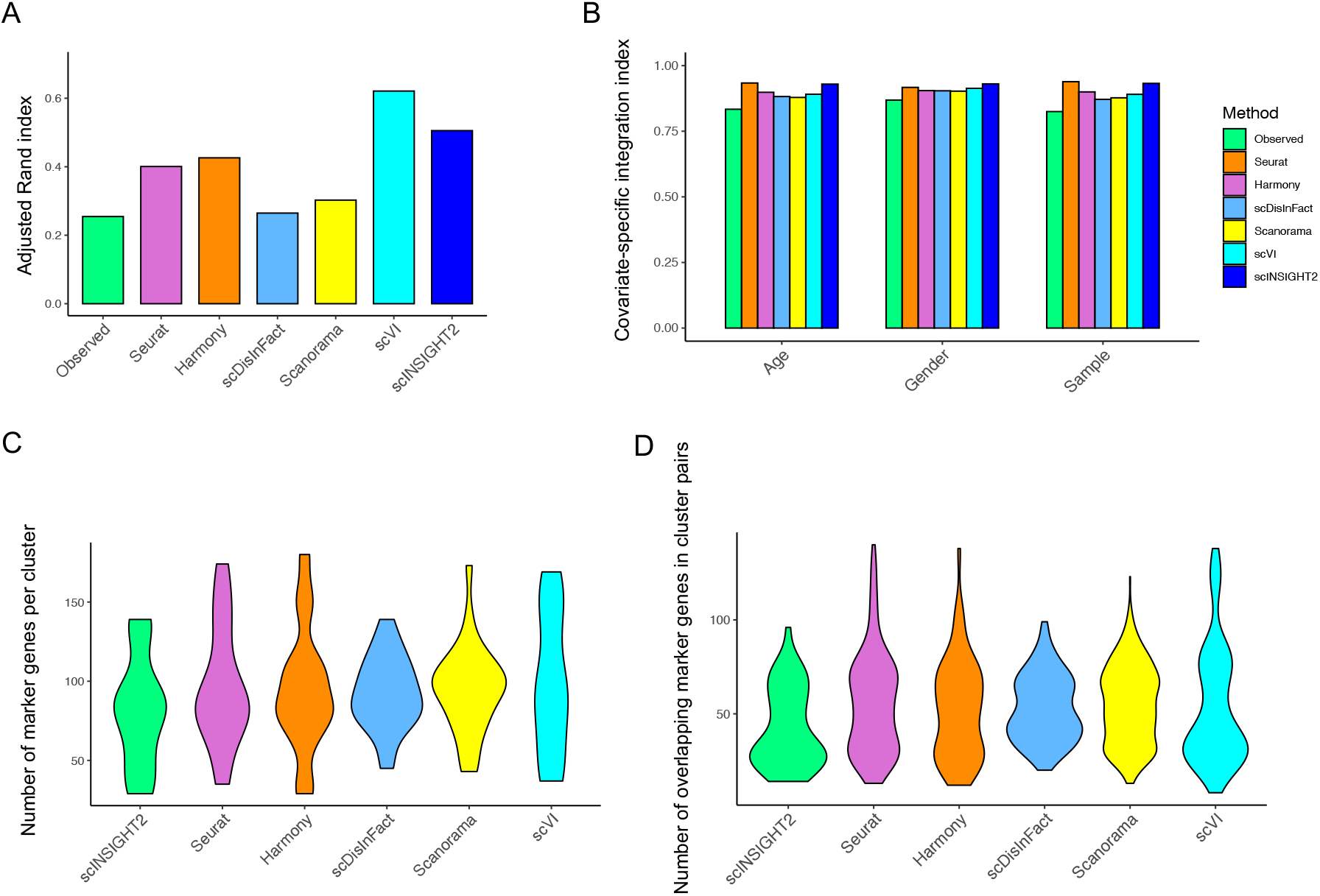
Application of integration methods to pancreatic single-cell samples. (**A**) Adjusted Rand index of cell clusters identified from observed and integrated data. (**B**) Covariate-specific integration index of the observed and integrated data by different methods. (**C**) Violin plots of marker gene number identified in cell clusters inferred by different methods. (**D**) Violin plots of the number of overlapping marker genes between pairs of clusters inferred by different methods.

To analyze if the difference in ARI score between scVI and scINSIGHT2 was due to cell subtypes that were not reflected in the annotated cell type labels, we focused on one cell type, the acinar cells. Acinar cells play a critical role in the production and secretion of digestive enzymes, and encompassed several clusters (mostly C2, C4, C6, and C8) identified by scINSIGHT2 (Supplementary Figure S6). It was evident from the tSNE plots that these clusters were characterized by different expression patterns of metagenes discovered by scINSIGHT2. Specifically, metagene 5 was highly expressed in clusters C6 and C8; metagene 8 was highly expressed in cluster C4; metagene 15 was highly expressed in clusters C2, C4, and C8. Given that clusters C2, C4, and C8 identified by scINSIGHT2 represented the largest subpopulations of acinar cells, we performed differential gene expression analysis focused on these three groups (see Methods). We found 71, 60, and 71 marker genes in clusters C2, C4 and C8 respectively. Supplementary Figure S7 displays the top three marker genes with the lowest adjusted *P* -values that were upregulated in clusters C2, C4 and C8, respectively. This provided further evidence that these clusters may represent distinct subtypes of acinar cells.

Next, we performed differential gene expression analysis to identify marker genes of clusters generated by different methods. The number of marker genes in scINSIGHT2 clusters ranged between 29 and 139 (Figure 3C). We also observed that the variation in marker gene number was larger for Seurat, Harmony, scVI, and Scanorama compared with scINSIGHT2 and scDisInFact, although the average numbers were similar across methods. We then calculated the overlap in marker genes between every pair of clusters. Ideally, if the clusters were truly distinct—representing biologically different cell populations—they should exhibit minimal overlap in their marker genes. Overall, clusters identified by scINSIGHT2 presented fewer overlapping marker genes compared to those identified by other methods (Figure 3D).

### 3.3 Application to COVID-19 data

To evaluate the performance of scINSIGHT2 on data encompassing different disease conditions, we applied it to a COVID-19 dataset with single-cell samples from seven COVID-19 patients and six healthy donors [31]. After preprocessing, the dataset comprised 44, 721 cells representing 13 distinct cell types. Additionally, available individual-level covariate information included age, gender, and disease condition.

On this dataset, scINSIGHT2 identified a total of 17 clusters (Figure 4A), with the corresponding cell type annotations from the original publication presented in Figure 4B. The metagenes identified by scINSIGHT2 exhibited distinct expression patterns across the clusters (Supplementary Figure S8). For example, metagenes 3, 4, 5, and 7 were highly expressed in red blood cells, plasma B cells, B cells, and CD14 monocytes, respectively. We compared our clustering results with those obtained using Seurat, Harmony, scDisInFact, Scanorama, and scVI, evaluating performance based on the ARI scores and the covariate-specific integration indices (Figures 4C-D, Supplementary Figure S9). scINSIGHT2 achieved the highest ARI score of 0.611, followed closely by scVI with a score of 0.605, while other methods yielded ARI scores below 0.5. Upon evaluating the covariate-specific integration indices, we observed that cluster C1 in scINSIGHT2’s results was distinctly characterized by age and sample ID (Supplementary Figure S10). Further investigation revealed that this cluster specifically represented red blood cells, which were unique to sample C6. This observation demonstrates scINSIGHT2’s capability to preserve rare cell types not commonly shared across all single-cell samples during the integration process.

**Figure 4:**
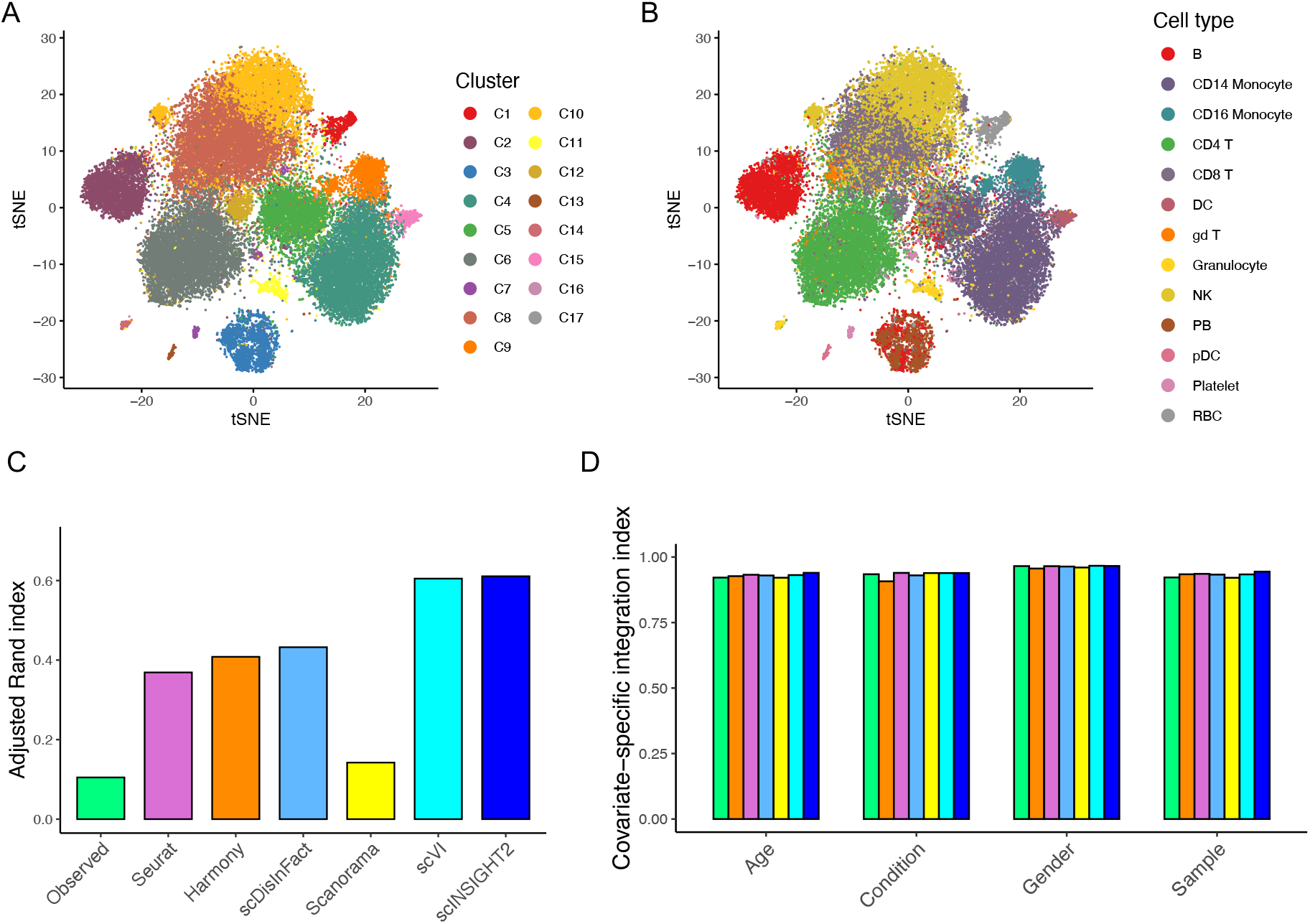
Comparison of observed and integrated data in the analysis of COVID-19 samples. (**A**) tSNE plot colored by scINSIGHT2-inferred cell clusters. (**B**) tSNE plot colored by annotated cell types.(**C**) Adjusted Rand index of cell clusters identified from observed and integrated data. (**D**) Covariate-specific integration index of the observed and integrated data.

Based on scINSIGHT2’s results, we compared the proportions of the identified cell clusters between the COVID-19 patients and healthy donors. For each cluster, we applied a logistic regression model to assess whether there was a significant association between the cell cluster proportion and the disease condition, while controlling for age and gender. This analysis identified clusters C4 and C10 as the most significantly associated with the disease condition, with cluster C4 enriched in COVID-19 samples and cluster C10 enriched in healthy samples (Figure 5A). We identified 85 DEGs between these two clusters (see Methods). Gene ontology (GO) enrichment analysis on genes up-regulated in cluster C4 revealed multiple enriched GO terms related to major histocompatibility complex (MHC) binding (Figure 5B), which was reported to be suppressed both transcriptionally and functionally by SARS-CoV-2 [36, 37]. We also conducted the same analysis on the two most significant clusters inferred by other integration methods (Figure 5C). scINSIGHT2 and Harmony led to the largest numbers of DEGs, indicating significant biological difference between condition-associated clusters. To assess the robustness of clustering results from integrated data, we identified DEGs between the two most significant clusters for each method, but solely using cells from individual samples. Figure 5D illustrates the overlap between DEGs identified from individual samples and those from all available samples. scINSIGHT2 and Harmony exhibited the highest DEG overlap, demonstrating their effectiveness in robustly distinguishing clusters associated with different conditions.

**Figure 5:**
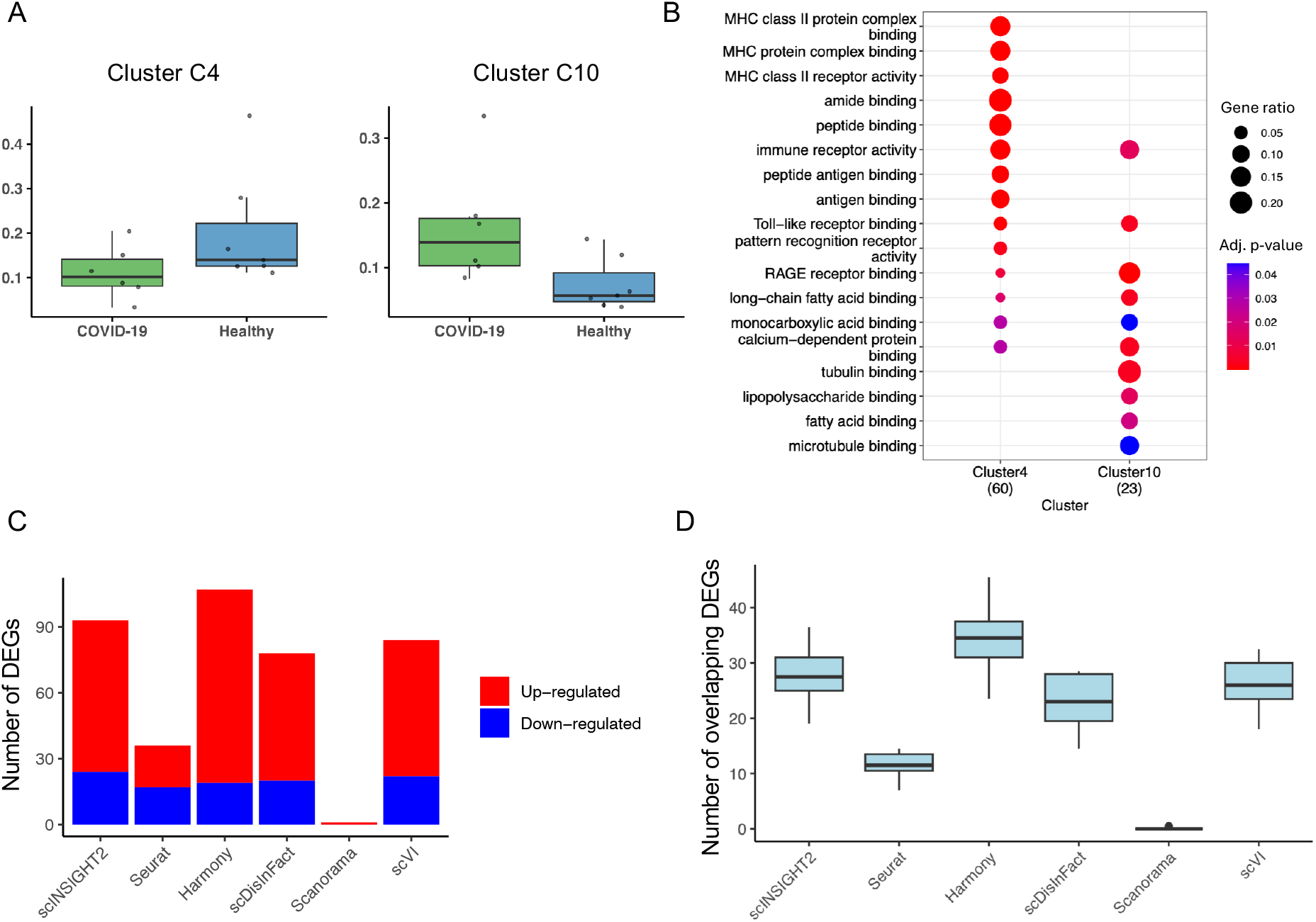
Comparison between condition-associated clusters in COVID-19 analysis. (**A**) Proportion of cells belonging to cluster C4 and cluster C10 in different COVID-19 and healthy samples. (**B**) Top enriched GO terms in genes upregulated in cluster C4 and cluster C10. (**C**) Total number of DEGs identified between the two most significant clusters from different methods. Up-regulated (or down-regulated) genes refer to genes with significantly higher expression in clusters enriched in COVID-19 (or healthy) samples. (**D**) Number of DEGs identified from individual samples that overlapped with the DEGs identified from all samples together.

## 4 Discussion

In this work, we introduce a new integration model named scINSIGHT2 for harmonizing gene expression data from multiple single-cell samples. scINSIGHT2 takes as input a concatenated count matrix from multiple samples across various biological groups, along with associated biological covariates. The biological covariates can include both discrete variables, such as disease status, and continuous variables, such as age. The model outputs inferred cell embeddings based on metagenes, which capture cellular identities and are used for clustering analysis and cell type annotation. In both simulation studies and real data analyses, scINSIGHT2 consistently demonstrated high accuracy and robustness.

We summarized the running time and memory usage of scINSIGHT2, Seurat, Harmony, scVI, Scanorama, and scDisInFact in Supplementary Table S2. As scINSIGHT2 incorporates individual-level information and solves a generalized matrix factorization optimization problem, it requires more computational resources than alternative methods. We recommend that users run the model with different numbers of metagenes or initializations in parallel to optimize performance.

In real data applications, identifying the specific subject-level covariates that contribute to gene expression differences between subjects can be challenging. We recommend including all available covariate information in the scINSIGHT2 model, allowing the model to infer the effects of these covariates directly from the data. If a covariate has little or no association with gene expression, the corresponding regression coefficients in the GLLVM will approach zero, minimizing its influence on the integration results. This robustness is demonstrated in our simulation study, where the inclusion of a covariate with no effect showed negligible impact on the model’s performance. Our current model considers only the linear effects of subject-level covariates, prioritizing simplicity and interpretability. However, if additional data provides evidence of non-linear relationships for specific covariates, the model could be extended to accommodate more complex patterns.

scINSIGHT2 assumes that the effects of biological covariates are additive to the latent factors corresponding to metagenes. This additive framework enables the straightforward incorporation of diverse covariates, enhancing computational efficiency and simplifying implementation. However, this assumption limits scINSIGHT2’s capacity to model higher-order relationships or interactions between covariates and latent factors. A promising future direction involves extending the model to account for more complex interactions. Recent approaches like scANVI [38] and STACAS [39] have employed supervised or semi-supervised integration strategies to achieve both data integration and cell type annotation simultaneously. Incorporating similar strategies into scINSIGHT2 could further enhance its utility. As cohort studies generating scRNA-seq data continue to grow, the availability of training data with subject-level covariates will provide robust support for such extensions.

## 5 Data Availability

The first pancreatic dataset was downloaded from GEO with access number GSE84133. The second pancreatic dataset was downloaded from https://doi.org/10.6084/m9.figshare.12420968. v8. The COVID-19 dataset was downloaded from https://cellxgene.cziscience.com/collections/a72afd53-ab92-4511-88da-252fb0e26b9a.

## 6 Acknowledgement

We would like to express our sincere gratitude to Shiwei Fu and other members of the Vivian Li Lab, as well as the Wenxiu Ma Lab, for their valuable discussions and insights throughout the course of this project. This work was partially supported by NIH R35GM142702. The authors acknowledge the HPC Center at UC Riverside (HPCC) and NSF-MRI grant 2215705 for the computing resources made available for conducting the research reported in this paper.

