## Supplementary Materials for "Harmonizing Heterogeneous Single-cell Gene Expression Data with Individual-level Covariate Information"

Yudi Mu[1] and Wei Vivian Li[1]

[1]Department of Statistics, University of California Riverside, Riverside, CA 92521, USA

### Supplementary Methods

#### Optimization procedure in scINSIGHT2

Given the objective function $Q(\Theta')$ derived in Methods, the gradient and Hessian matrix of $u_i$ are

$$\frac{\partial Q}{\partial u_i} = -\sum_{j=1}^{M} (y_{ij} - \mu_{ij}) \lambda_j + \frac{1}{\sigma^2} u_i,$$

$$\frac{\partial^2 Q}{\partial u_i \partial u_i^\top} = \sum_{j=1}^{M} \mu_{ij} \lambda_j \lambda_j^\top + \frac{1}{\sigma^2} I_P \quad \text{and} \quad \frac{\partial^2 Q}{\partial u_i \partial u_k^\top} = 0 \text{ for } k \neq i.$$

Similarly, for $\beta_j$ and $\lambda_j$, the gradients and Hessian matrices are

$$\frac{\partial Q}{\partial \beta_j} = -\sum_{i=1}^{N} (y_{ij} - \mu_{ij}) x_i,$$

$$\frac{\partial Q}{\partial \lambda_j} = -\sum_{i=1}^{N} (y_{ij} - \mu_{ij}) u_i,$$

and

$$\frac{\partial^2 Q}{\partial \beta_j \partial \beta_j^\top} = \sum_{j=1}^{N} \mu_{ij} x_i x_i^\top,$$

$$\frac{\partial^2 Q}{\partial \lambda_j \partial \lambda_j^\top} = \sum_{j=1}^{N} \mu_{ij} u_i u_i^\top,$$

$$\frac{\partial^2 Q}{\partial \beta_j \partial \lambda_j^\top} = \sum_{j=1}^{N} \mu_{ij} x_i u_i^\top,$$

$$\frac{\partial^2 Q}{\partial \beta_j \partial \beta_k^\top} = \frac{\partial^2 Q}{\partial \lambda_j \partial \lambda_k^\top} = \frac{\partial^2 Q}{\partial \beta_j \partial \lambda_k^\top} = 0 \text{ for } k \neq j.$$

In addition, to update the standard deviation $\sigma$, we have

$$\frac{\partial Q}{\partial \sigma} = -\frac{\sum_{i=1}^N u_i^\top u_i}{\sigma^3} + \frac{NP}{\sigma},$$

$$\frac{\partial^2 Q}{\partial \sigma^2} = \frac{3\sum_{i=1}^N u_i^\top u_i}{\sigma^4} - \frac{NP}{\sigma^2}.$$

With the derived gradients and Hessian matrices, the estimation process can be summarized as the following steps:

(1) Initialize $U, B, \Lambda$ randomly, where $B = (\beta_1, \beta_2, \ldots, \beta_m) \in \mathbb{R}^D$. Randomly sample $\beta_j \sim N(0, 0.1 \times I_M)$, $u_i \sim N(0, 0.1 \times I_P)$, $\lambda_j \sim N(0, 0.1 \times I_P)$ $(i = 1, ..., N, j = 1, ..., M)$. Initialize $\sigma = \frac{\sum_{i=1}^N \text{SD}(u_i)}{N}$.

(2) Repeat the following steps to perform updates until the stopping rule is satisfied or maximum iteration number (default to 5000) is reached.

$$\begin{aligned}
U_{ik}^{(t+1)} &= u_i^{(t)} - \alpha \left[\nabla_\theta^2 Q\left(u_i^{(t)}\right)\right]^{-1} \nabla_\theta Q\left(u_i^{(t)}\right), \\
\lambda_j^{(t+1)} &= \lambda_j^{(t)} - \alpha \left[\nabla_\theta^2 Q\left(\lambda_j^{(t)}\right)\right]^{-1} \nabla_\theta Q\left(\lambda_j^{(t)}\right), \\
\beta_j^{(t+1)} &= \beta_j^{(t)} - \alpha \left[\nabla_\theta^2 Q\left(\beta_j^{(t)}\right)\right]^{-1} \nabla_\theta Q\left(\beta_j^{(t)}\right), \\
\sigma^{(t+1)} &= \sigma^{(t)} - \alpha\left(\frac{\partial^2 Q}{\partial(\sigma^{(t)})^2}\right)^{-1} \frac{\partial Q}{\partial \sigma^{(t)}}
\end{aligned}$$

(3) We compare the estimated cell embeddings at two adjacent steps using $\text{RMSE}_U(t) \triangleq \frac{1}{NP}\left(\sum_{i=1}^N \sum_{p=1}^P (U_{ip}^{(t+1)} - U_{ip}^{(t)})^2\right)^{\frac{1}{2}}$. The estimation is considered converged if $\text{RMSE}_U(t) < \epsilon_1$ and the average RMSE of the previous 100 iterations $\frac{1}{100}\sum_{q=t-99}^t \text{RMSE}_U(q) < \epsilon_2$, where $\epsilon_1$ and $\epsilon_2$ are sufficiently small values.

(4) Perform a rotation on the final estimates $\hat{U}$ and $\hat{\Lambda}$ for parameter identifiability. Denote the covariance matrix of $\hat{U}$ as $S$. We first obtain a matrix $W$ such that $W^\top W = S^{-1}$, using the Cholesky algorithm. Then, we perform the following transformations: $\hat{U} \leftarrow \hat{U}W$ and $\hat{\Lambda} \leftarrow W^{-1}\hat{\Lambda}$. Then, we find a QR decomposition of $\hat{\Lambda}$ such that $\hat{\Lambda} = QR$, and update $\hat{U}$ and $\hat{\Lambda}$: $\hat{U} \leftarrow \hat{U}Q$ and $\hat{\Lambda} \leftarrow R$. This step ensures that $\hat{\Lambda}$ is a lower triangular matrix. Denote vector $d = \text{diag}(\hat{\Lambda})$, vector $s = \text{sign}(d)$, we get $\hat{U} \leftarrow (s\hat{U}^\top)^\top$, $\hat{\Lambda} \leftarrow s\hat{\Lambda}$. This will make $\hat{\Lambda}$ have positive diagonals.

### Normalization of estimated cell embeddings

Since our estimation process relies on the Laplace approximation and may not directly yield cell embeddings that are optimal for clustering, we implement a normalization procedure from scINSIGHT to further refine the cell embeddings before clustering analysis.

(1) We use $\hat{U}_1 \in \mathbb{R}_{n_1 \times M}, \ldots, \hat{U}_K \in \mathbb{R}_{n_K \times M}$ to represent the estimated cell embeddings of the $K$ samples. Each row of $\hat{U}_k$ $(k = 1, \ldots, K)$ is normalized so that its $L_2$ norm is $1$. We denote the normalized matrices by $\tilde{U}_k$.

(2) For each pair of two samples, we find $20$ nearest neighbours of each cell in one sample from the other sample based on $\tilde{U}_k$'s. Then, a mutual nearest neighbour graph of all cells is constructed based on the mutual nearest neighbours between every pair of samples.

(3) The Louvain method is used to perform clustering on the mutual nearest neighbour graph obtained in the previous step. Within each cluster, we perform the quantile normalization on $\tilde{U}_k$'s to reduce batch effects within clusters.

### Supplementary Figures

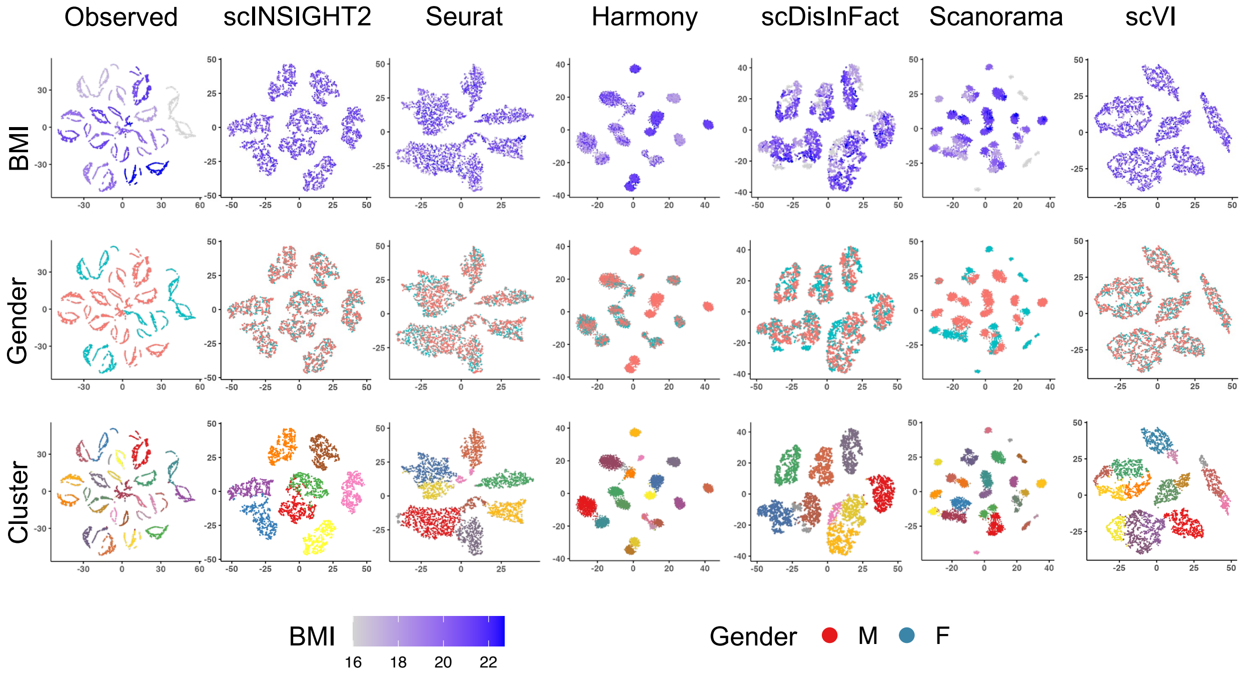


**Figure S1:** Comparison between integration methods in the simulation study (under first setting). For each method, tSNE plots with cells colored by BMI, gender, and inferred cluster are displayed.

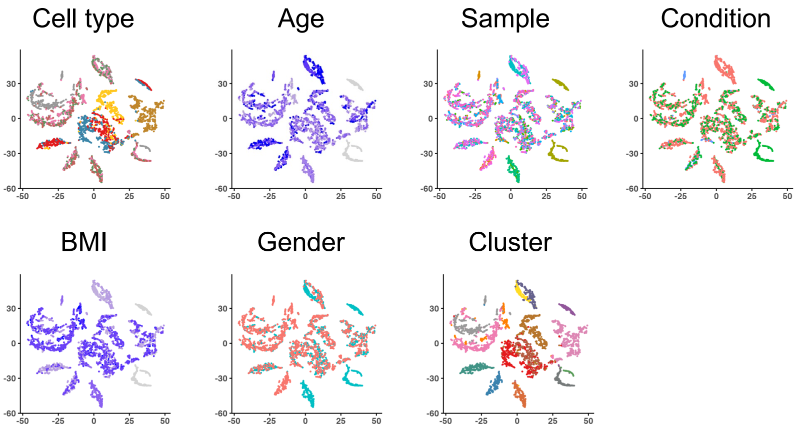


**Figure S2:** Integration result of scINSIGHT. The tSNE plots are colored by cell type, age, sample, condition, BMI, gender, and cluster.

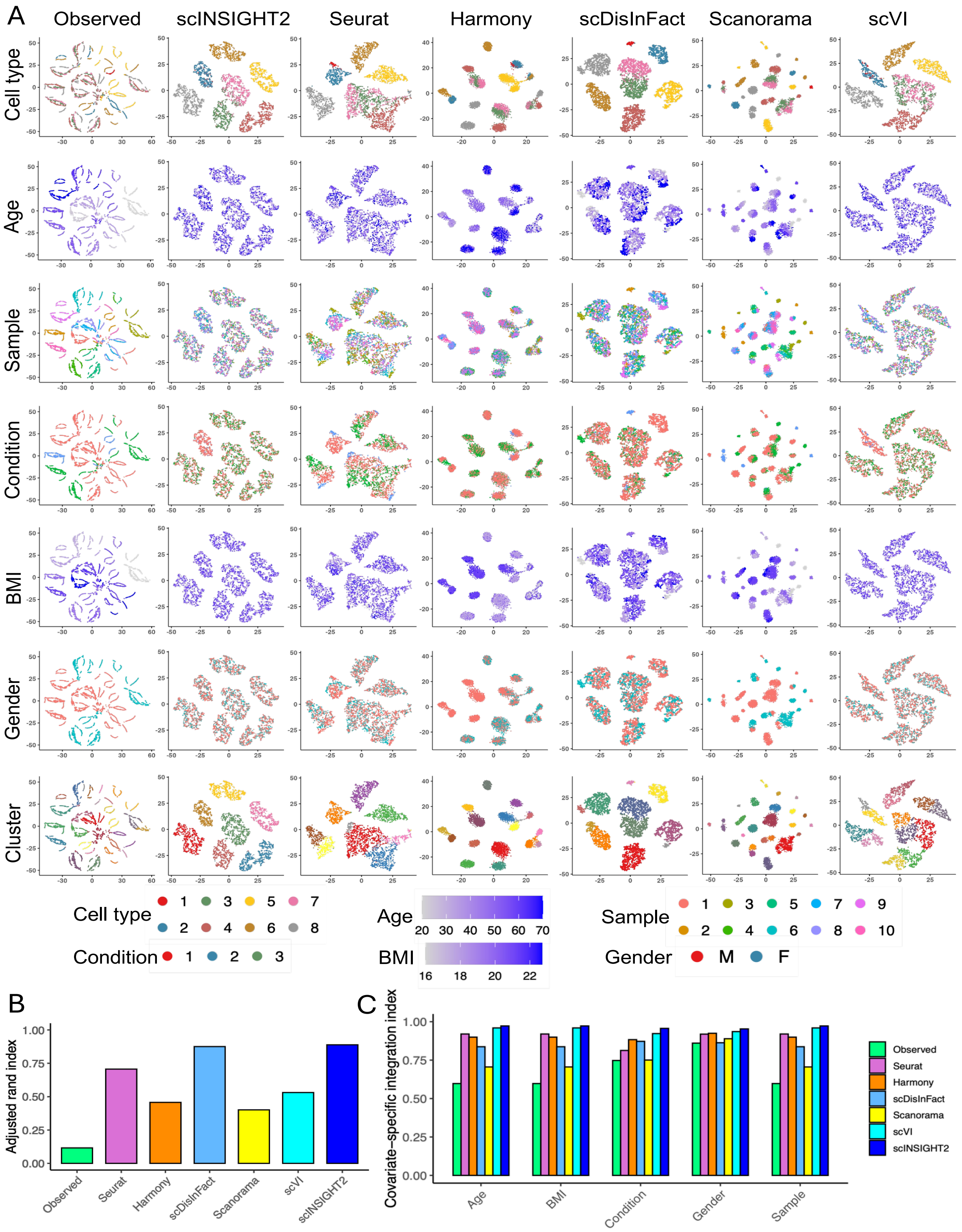


**Figure S3:** Comparison between integration methods in the simulation study with unique cell types. (**A**) tSNE plots of observed data and integrated data by different methods. For each method, tSNE plots colored by cell type, age, sample, condition, BMI, gender and cluster are displayed. (**B**) Adjusted Rand index calculated using clusters identified from the observed or integrated data. (**C**) Covariate-specific integration index of the observed and integrated data.

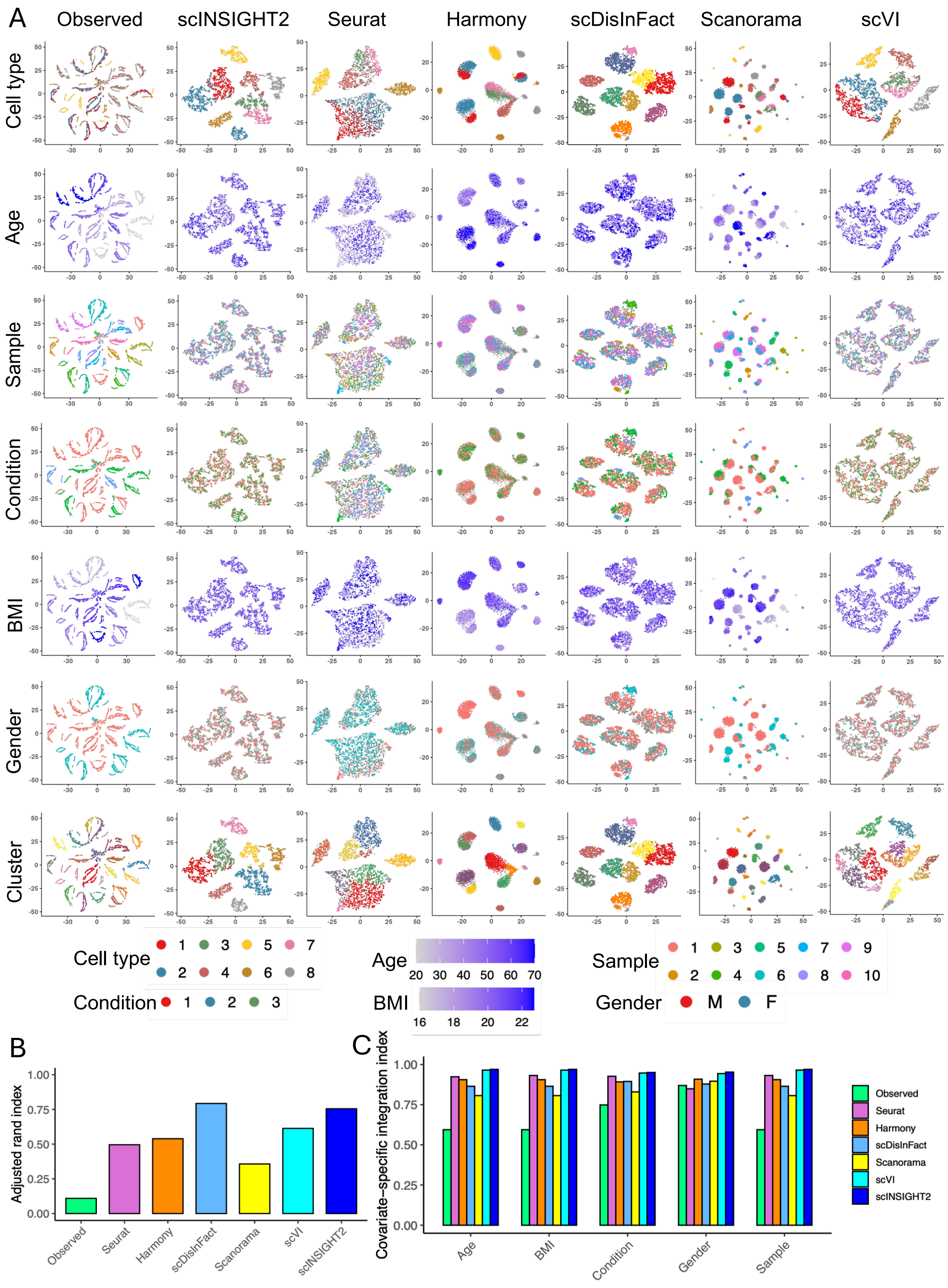


**Figure S4:** Comparison between observed data and different integration methods in the simulation study with age-associated cell types. (**A**) tSNE plots of observed data and integrated data by different methods. For each method, tSNE plots colored by cell type, age, sample, condition, BMI, gender and cluster are displayed. (**B**) Adjusted Rand index calculated using clusters identified from the observed or integrated data. (**C**) Covariate integration index of the observed and integrated data.

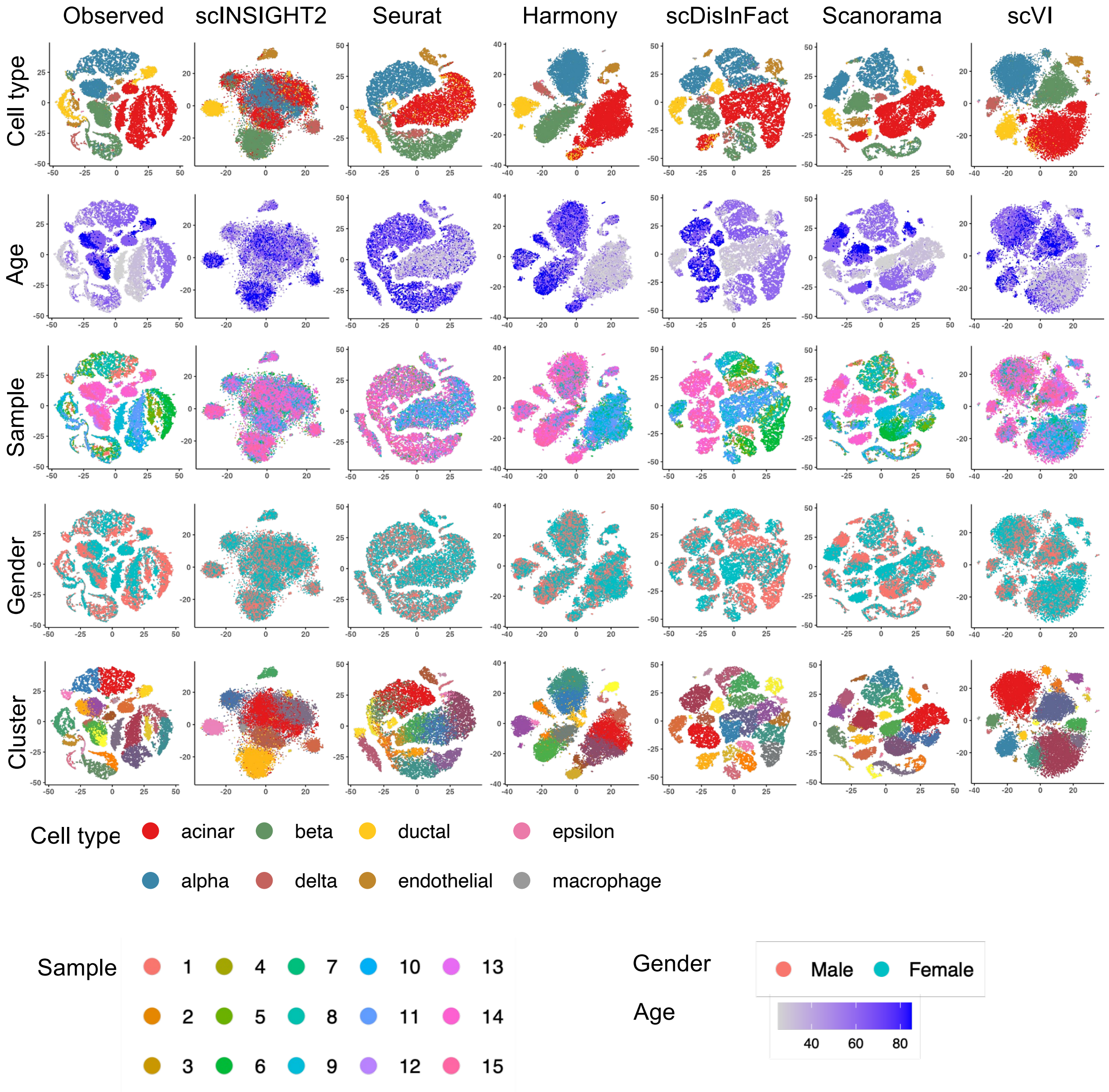


**Figure S5:** Comparison between observed data and integrated data by different integration methods in the pancreatic study. For each method, tSNE plots colored by cell type, age, sample, gender and cluster are displayed.

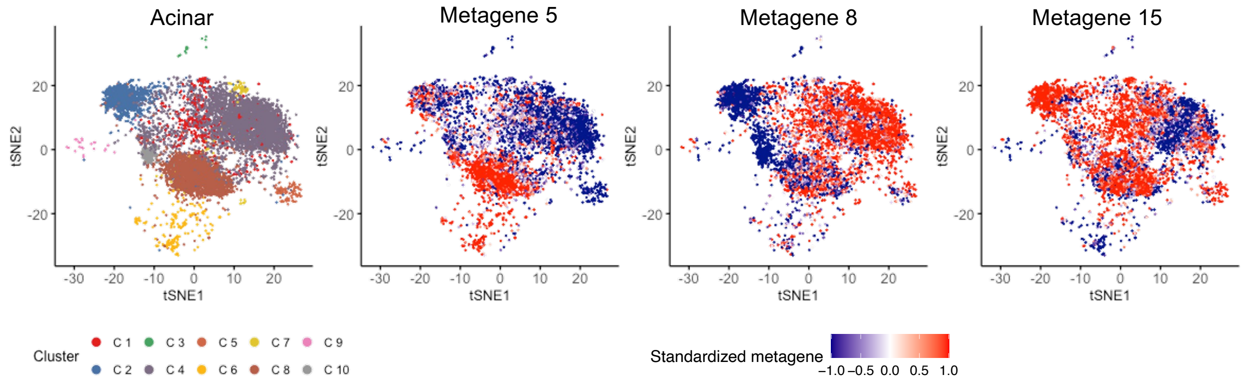


**Figure S6:** tSNE plots of acinar cells based on integrated data by scINSIGHT2. From left to right, the plots are colored by cluster and cell scores corresponding to metagene $5$, metagene $8$, and metagene $15$.

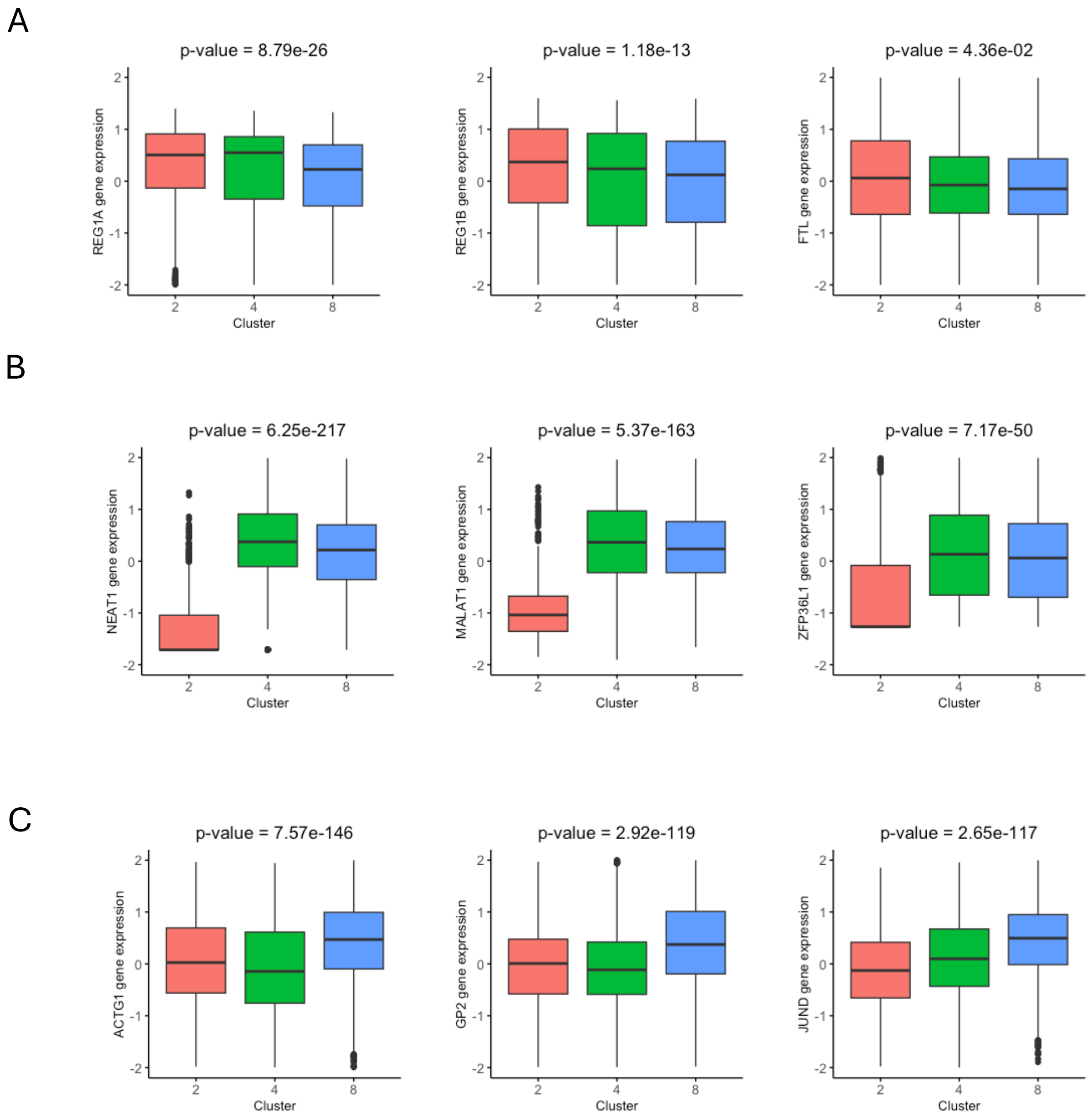


**Figure S7:** Top three marker genes of clusters C2, C4 and C8 in acinar cells. The $P$-values from Wilcoxon test are shown on the top of each figure. (**A**) Boxplots of top three marker genes upregulated in cluster C2. (**B**) Boxplots of top three marker genes upregulated in cluster C4. (**C**) Boxplots of top three marker genes upregulated in cluster C8.

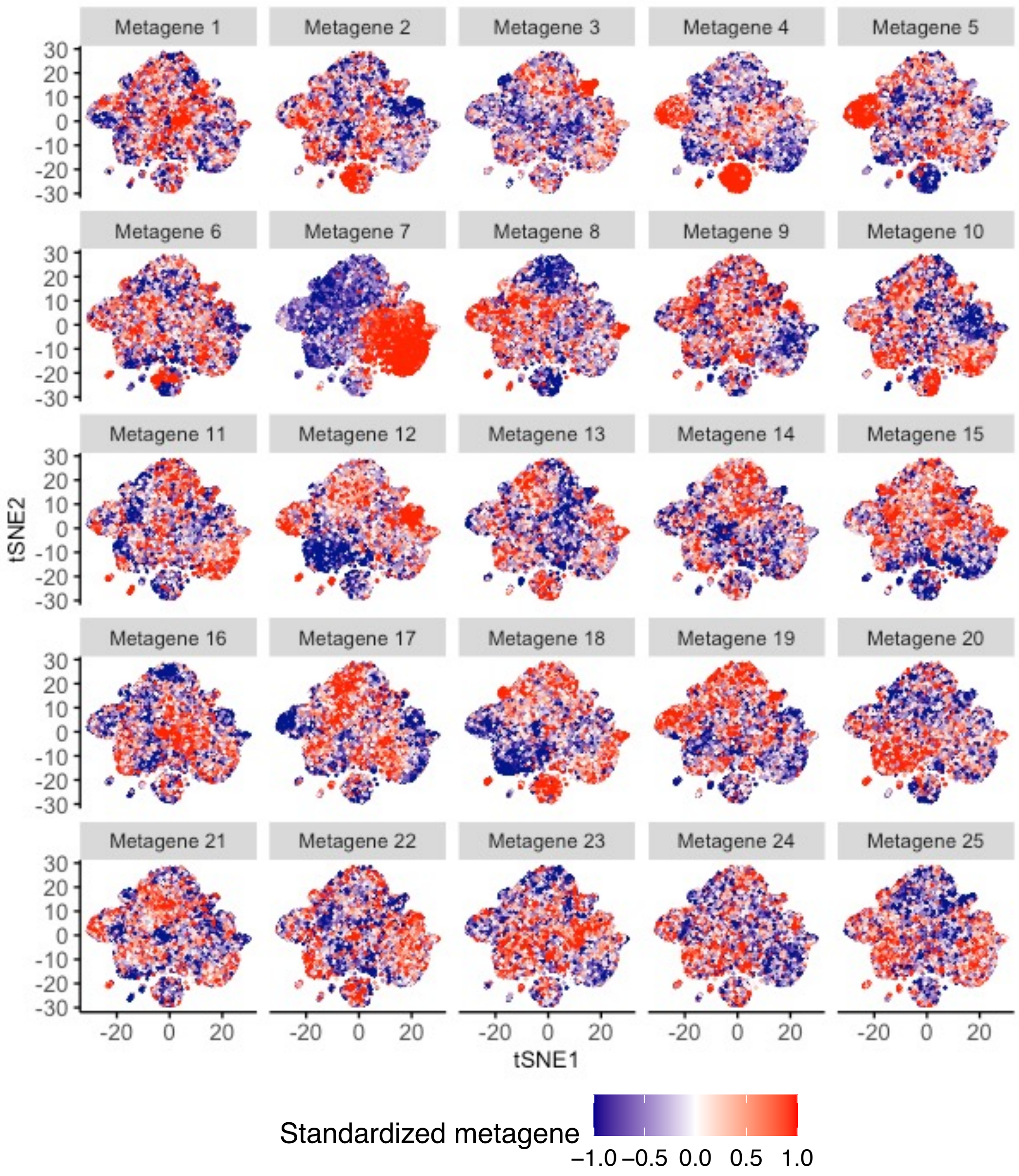


**Figure S8:** t-SNE plots based on integrated COVID-19 data processed by scINSIGHT2, with cells colored according to their normalized scores on metagenes.

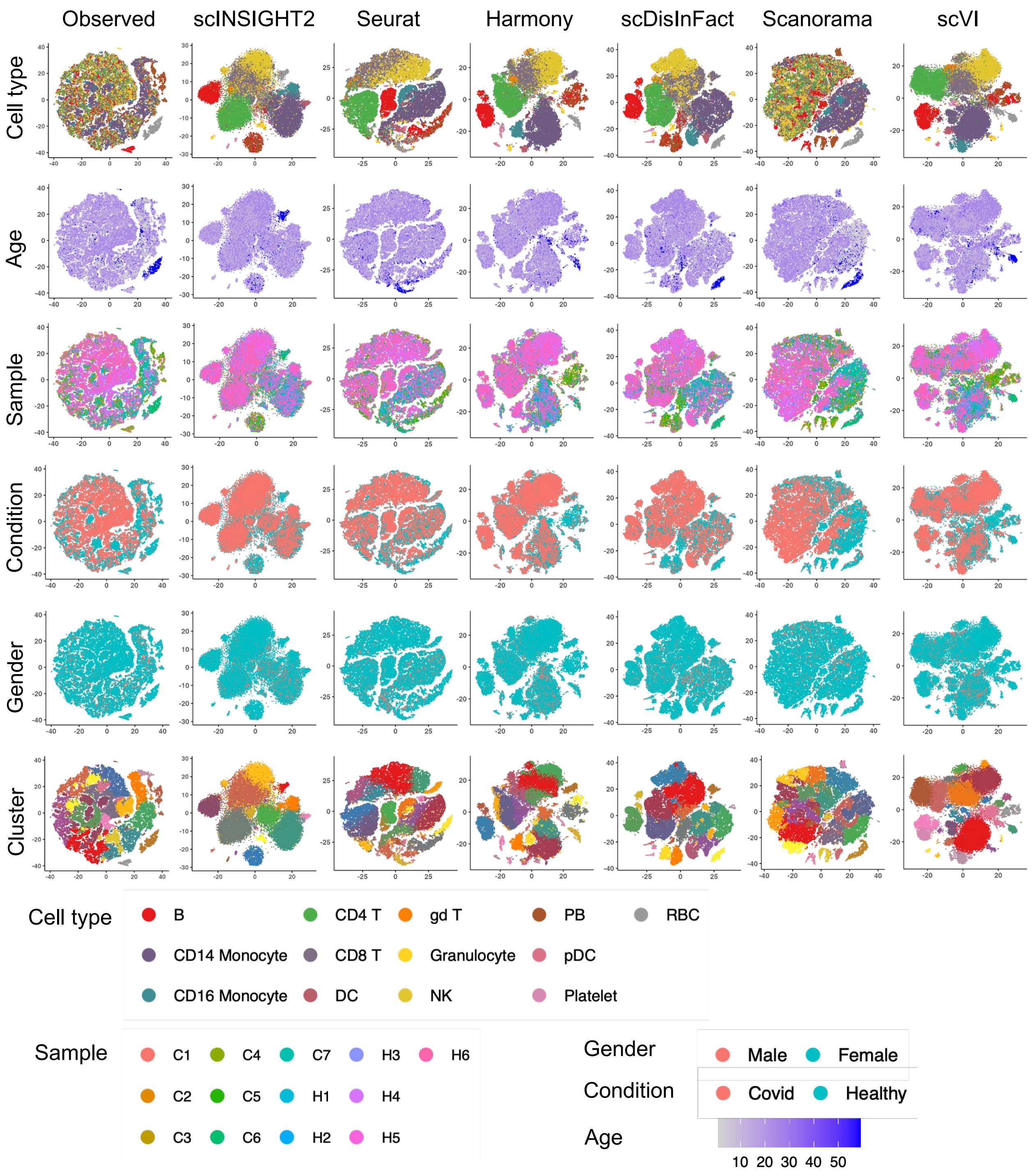


**Figure S9:** Comparison between observed data and different integration methods in the COVID-19 study. For each method, tSNE plots of observed data and integrated data colored by cell type, age, sample, condition, gender and cluster are displayed.

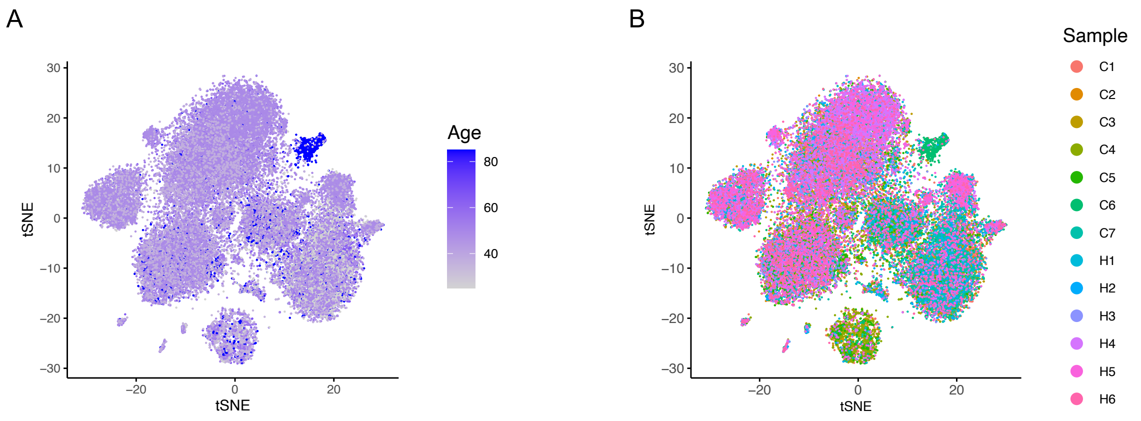


**Figure S10:** tSNE plots based on integrated COVID-19 data by scINSIGHT2, colored by age (**A**) or sample (**B**).

### Supplementary Tables

**Table S1:** Mean parameters of normal distribution for the simulation of cell embeddings.

| cell type | p=1 | p=2 | p=3 | p=4 | p=5 |
|---|---|---|---|---|---|
| type1 | $-1.2$ | $-0.5$ | $0$ | $0.5$ | $1.2$ |
| type2 | $-0.5$ | $0$ | $0$ | $0.5$ | $1.2$ |
| type3 | $1.2$ | $0.5$ | $0$ | $-0.5$ | $-1.2$ |
| type4 | $1.2$ | $0.5$ | $0$ | $-1.2$ | $-0.5$ |
| type5 | $-1.2$ | $0$ | $0.5$ | $1.2$ | $-0.5$ |
| type6 | $-1.2$ | $0$ | $0.5$ | $-1.2$ | $0.5$ |
| type7 | $1.2$ | $0.5$ | $0$ | $0$ | $-0.5$ |
| type8 | $0.5$ | $0.5$ | $0$ | $-0.5$ | $1.2$ |

**Table S2:** Computational time and memory usage of scINSIGHT2 and alternative methods. The time and memory usage reported for scINSIGHT2 correspond to the optimal number of metagenes and initialization seed.

| Dataset | scINSIGHT2 | Seurat | Harmony | Scanorama | scVI | scDisInFact |
|---|---|---|---|---|---|---|
| Simulation<br>$5,000$ cells<br>$10$ samples<br>$5$ covariates | $84$ mins<br>$2.01$ GB | $3.8$ mins<br>$1.4$ GB | $0.43$ mins<br>$0.919$ GB | $0.82$ mins<br>$0.367$ GB | $2.7$ mins<br>$2.2$ GB | $5.3$ mins<br>$0.746$ GB |
| Pancreas<br>$27,435$ cells<br>$15$ samples<br>$3$ covariates | $660$ mins<br>$9.56$ GB | $30$ mins<br>$3.26$ GB | $2.08$ mins<br>$1.537$ GB | $3.8$ mins<br>$3.32$ GB | $10.55$ mins<br>$3.63$ GB | $7.93$ mins<br>$0.88$ GB |
| COVID-19<br>$44,721$ cells<br>$13$ samples<br>$4$ covariates | $1680$ mins<br>$15.26$ GB | $71$ mins<br>$5.47$ GB | $3.54$ mins<br>$2.64$ GB | $36$ mins<br>$8.98$ GB | $11.6$ mins<br>$5.98$ GB | $12$ mins<br>$0.835$ GB |
